# Improved Spectral Inversion of Blood Oxygenation due to Reduced Tissue Scattering: Towards NIR-II Photoacoustic Imaging

**DOI:** 10.1101/2024.08.08.607178

**Authors:** Vinoin Devpaul Vincely, Carolyn L. Bayer

**Affiliations:** Department of Biomedical Engineering, Tulane University, New Orleans, LA 70112, USA

**Keywords:** spectral photoacoustic imaging, blood oxygenation, NIR-II, spectral coloring, in vivo

## Abstract

**Significance:** Conventional spectral photoacoustic imaging (sPAI) to assess tissue oxygenation (sO_2_) uses optical wavelengths in the first near infrared window (NIR-I). This limits the maximum imaging depth (∼1 cm) due to high spectral coloring of biological tissues.

**Aim:** Second near infrared or short-wave infrared (NIR-II or SWIR) wavelengths (950-1400 nm) show potential for deep tissue sPAI due to the exponentially reduced tissue scattering and higher maximum exposure threshold (MPE) in this wavelength range. However, to date, a systematic assessment of NIR-II wavelengths for sPAI of tissue sO_2_ has yet to be performed.

**Approach:** The NIR-II PA spectra of oxygenated and deoxygenated hemoglobin was first characterized using a phantom. Optimal wavelengths to minimize spectral coloring were identified. The resulting NIR-II PA imaging methods were then validated *in vivo* by measuring renal sO_2_ in adult female rats.

**Results:** sPAI of whole blood under a phantom and of circulating renal blood in vivo, demonstrated PA spectra proportional to wavelength-dependent optical absorption. NIR-II wavelengths had a ∼50% decrease in error of spectrally unmixed blood sO_2_ compared to conventional NIR-I wavelengths. In vivo measurements of renal sO_2_ validated these findings and demonstrated a ∼30% decrease in error of estimated renal sO_2_ when using NIR-II wavelengths for spectral unmixing in comparison to NIR-I wavelengths.

**Conclusions:** sPAI using NIR-II wavelengths improved the accuracy of tissue sO_2_ measurements. This is likely due to the overall reduced spectral coloring in this wavelength range. Combined with the increased safe skin exposure fluence limits in this wavelength range, demonstrate the potential to use NIR-II wavelengths for quantitative sPAI of sO_2_ from deep heterogeneous tissues.

## 1. Introduction

Oxygen saturation of hemoglobin (sO_2_) is an important physiological metric for clinical applications in oncology [1, 2], neurology [3–7], cardiology [8], and obstetrics [9–11]. Spectral photoacoustic imaging (sPAI) is a promising imaging modality to assess deep tissue sO_2_ due to its relatively low cost and real-time imaging capability. The advantages of PAI result from the use of optical excitation to generate broadband acoustic waves through a phenomenon called thermoelastic expansion – an instantaneous conversion of optical to mechanical energy when chromophores absorb light. In contrast to purely optical techniques, PAI allows visualization of deeper structures, due to the reduced sound attenuation by biological tissues.

Photoacoustic (PA) image contrast is directly proportional to the local optical fluence and the concentration of optical chromophores, including oxygenated (oxy-Hb) and deoxygenated (deoxy-Hb) hemoglobin, within the imaged region. sPAI leverages the distinct optical spectral features of oxy-Hb & deoxy-Hb by acquiring multiple images at different optical wavelengths to estimate their spatially varying concentration within the imaged tissue, typically through linear spectral unmixing of the resulting photoacoustic signals. One of the earliest demonstrations of measuring blood sO_2_ using spectral PA imaging was performed by Wang et. al. to monitor cerebral vasculature in small-animal models, with imaging performed between 580-600 nm [12]. sPAI has been extended to a variety of biomedical applications - Diot et. al. performed blood sO_2_ up to depths of 2.2 cm in breasts containing invasive lobular carcinoma using excitations between 700- 970 nm [13]. More recently, our group measured placental oxygenation in normal pregnant and preeclamptic rats with optical excitations at 690, 808 & 950 nm, up to a depth of 2 cm [10].

The accuracy of sO_2_ estimated using sPAI can be affected by the wavelength- & spatial-variant local fluence at the imaged target. Photons undergo a complex wavelength-dependent attenuation as they propagate through heterogenous biological tissues due to the spectrally and spatially varying optical properties (absorption and scattering) of different tissue types. This results in distortions in the measured PA spectrum of the imaged targets, called spectral coloring. Various studies have attempted to account for the spectrally varying local fluence [14–19]. Optical transport models, such as Monte-Carlo (MC) simulations, have been widely used to model the light distribution in complex biological tissues to account for the spatially varying local fluence and improve the accuracy of blood oxygenation estimations [14–17]. However, accurate optical transport models require *a priori* knowledge of the optical properties of the heterogenous tissues with the imaged plane. Furthermore, MC models require powerful computational hardware/time to accurately simulate fluence maps, making clinical translation for real time imaging difficult. Direct measurements of local fluence using diffuse optical tomography (DOT), have also been demonstrated [18, 19]. However, in addition to requiring additional imaging hardware, DOT has insufficient spatial accuracy to adequately correct for tissue optical properties.

Conventionally, sPAI for the measurement of sO_2_ has been implemented using wavelengths in the NIR-I window (690-950 nm) [9, 10, 12]. However, high spectral coloring within this optical window limits quantitative measurements at larger in vivo depths (beyond ∼1 cm). sPAI using short-wave infrared (SWIR) or second near infrared (NIR-II) wavelengths (950-1800 nm) offers advantages including exponentially reduced optical scattering by biological tissues, higher maximum safety exposure limits (MPE) for skin, and reduced auto-fluorescence [20], potentially facilitating deep tissue sPAI of sO_2_. Deep in vivo sPAI using NIR-II wavelengths has been challenged due to exponentially increasing water absorption that limits maximum light penetration in tissues. Therefore, to date, NIR-II PA imaging has typically been limited to the use of 1064 nm for the anatomical visualization of vasculature [4, 21, 22], demonstrating imaging at in vivo depths up to 4 cm [4]. However, the use of the NIR-II window for sPAI-derived measurements of sO_2_ has yet to be demonstrated. Recently, Carr et. al. demonstrated high contrast deep fluorescence imaging of in vivo mouse brain with an intact skin using 1450 nm (water absorption peak) [23]. They attributed their high contrast at 1450 nm to a preferentially suppressed photon path length which decreases noise due to multiple scattering events and increases target absorption. This behavior of photons shows promise for deeper PA imaging using NIR-II wavelengths.

In this study, NIR-II characterization of hemoglobin PA spectra at distinct oxygenation levels was performed in a simple phantom. PA spectra of oxy-Hb and deoxy-Hb was observed to be consistent with their respective optical absorption. Spectral unmixing of whole blood sO_2_ using NIR-II wavelengths showed a ∼50% decrease in absolute error when compared to conventional NIR-I wavelengths. Next, sPAI of renal oxygenation using NIR-II wavelengths was performed. *In vivo* measurements of renal oxygenation validated phantom experiments by demonstrating a ∼30% reduction in absolute error when using NIR-II excitations for spectral unmixing in comparison to conventional NIR-I wavelengths.

## 2. Methods

### 2.1 Photoacoustic *imaging instrumentation*

PA imaging was performed using a pulsed optical parametric oscillator (OPO) laser (Phocus, BENCHTOP, Opotek Inc., Carlsbad, CA, USA) integrated with an open architecture data acquisition system (Vantage 256, Verasonics Inc., Kirkland, WA, USA), using a 6 MHz linear array transducer (L7-4, Philips, Amsterdam, Netherlands). The laser generates 5 ns pulses at a 10 Hz repetition rate at tunable excitation wavelengths in the NIR-I (690-950 nm) and NIR-II (1064 nm, 1200 nm-2400 nm) windows at 1 nm increments. The optical energy density incident on the surface of the imaged target was measured using a power meter (Ophir Technologies, West North Logan, UT, USA) and maintained with an average pulse-to-pulse variation in laser output of less than 5%. The illumination geometry of the beam was monitored using burn paper. All data analysis was performed in MATLAB.

### 2.2 Porcine phantom for NIR-II blood characterization

A freshly cut slice of porcine tissue with thicknesses of 3 mm was placed over a polytetrafluoroethylene (PTFE) tube (1.5/2.0 mm inner/outer diameter) embedded in a gelatin mold. This provided an optical path length of 0.3 cm. Defibrinated whole bovine blood (910-100, Quad Five, Ryegate, MT, USA) at two levels of oxygenation (sO_2_) – 100% and 0%, and a phosphate-buffer (PBS) solution were loaded in the tube sequentially. Deoxygenation of the stock hemoglobin was performed using sodium hydrosulfite (Na_2_S_2_O_4,_ 157953-100G, Sigma-Aldrich, St. Louis, MO, USA) at concentrations where conversion to methemoglobin is limited [24]. The stability of the targeted sO_2_ was monitored using a bare-fiber oxygen sensor coupled with an OxyLite system (Oxford Optronix, Oxford, United Kingdom) at regular intervals throughout imaging. PA images were collected at NIR-I (690 – 950 nm at 20 nm increments) and NIR-II (1064, 1200 – 1400 nm at 10 nm increments) at a 25 mJ/cm^2^ surface fluence (circular beam with diameter 0.5 cm). The phantom experiments were repeated 3 times (i.e. three separately prepared phantoms).

In the collected PA images, a region of interest (ROI) was selected around the tube and the pixels within this region averaged. To account for measurement variations due to tissue inhomogeneity across different experimental runs, all PA spectra measured for oxy-Hb and deoxy-Hb were normalized to the isosbestic point of hemoglobin (808 nm). Next, the local fluence at the tube within the phantom was calculated using Beer’s law by setting the attenuation coefficient of the porcine tissue to a combination of water and lipid absorption (the attenuation of the individual components of the porcine tissue is shown in the supplemental figure S1). The fluence at the illumination surface was measured using the power meter, and then used to normalize the raw PA signals to obtain the final PA spectra of oxy-Hb and deoxy-Hb.

### In vivo NIR-II imaging of renal vasculature in rat

All animal imaging and surgical procedures were performed following protocols approved by the Institutional Animal Care and Use Committee (IACUC) at Tulane University. Adult female Sprague Dawley rats (n = 3) were acquired from a commercial vendor (Charles River Laboratories, Boston, MA). The animals were anesthetized using 3% isoflurane and the hair on the abdomen and back was removed using a depilatory cream. The animal was then moved to a transparent sheet (polyethylene terephthalate, PET) for illumination of the posterior side of the animal. A circular beam of 1.3 cm diameter was used to deliver an optical fluence of 5-6 mJ/cm^2^ at the surface of the animal’s skin (accounting for any energy attenuation by the transparent sheet, Fig. S2). The breathing rate was monitored and the temperature was measured at the animal’s skin (LaserPro LP300, KIZEN) during the imaging session. The 6 MHz linear transducer was placed on the abdomen and positioned over the kidney of the animal. PA images were acquired at NIR-I (690 – 950 nm with 20 nm increments) and NIR-II (1064, 1200 – 1300 nm with 10 nm increments). After the imaging session, the oxygenation of the kidney was measured over 10 minutes using a surgically inserted oxygen probe. The animal was euthanized using excess CO_2_ for a total of 15- 20 minutes, followed again by PA imaging. The renal oxygenation of the euthanized rat was again measured after imaging, ∼1.5h post-euthanasia, using the surgically inserted oxygen probe.

To demonstrate that the measured PA signal was primarily generated by hemoglobin, the kidney of a euthanized animal was removed and bleached of hemoglobin by suspending the tissue in excess hydrogen peroxide overnight (∼ 15h) [25, 26]. The bleached kidney was then placed back in the abdomen of the euthanized animal and imaged. To characterize the PA spectra of the renal vasculature under different conditions (pre- and post-euthanasia, and bleached of hemoglobin), a ROI was defined around the kidney and the pixels within the region were averaged (Fig. 2c). To account for PA signal variation due to motion artifacts and variations in animal size, the spectral PA images were normalized at the isosbestic point (808 nm). Light attenuation due to the tissue absorption was calculated using Beer’s law. Light propagated through a layer of subcutaneous fat prior to illuminating the kidney (as seen in the inset of fig. 2a), therefore this light attenuation correction assumed a tissue composition primarily of water and lipids. This attenuation was then used to normalize the mean ROI PA signals to obtain a fluence normalized PA spectra of oxy-Hb and deoxy-Hb kidneys. Finally, a temperature difference between pre- (∼36 °C) and post- (∼28-30 °C) euthanasia was observed. Finally, A normalization factor derived as a ratio of pre- to post-euthanasia body temperature was used to adjust the PA spectra of the deoxy-Hb kidney in the NIR-II window (a schematic representation of the PA correction is outlined under fig. S6).

**Fig. 1:**
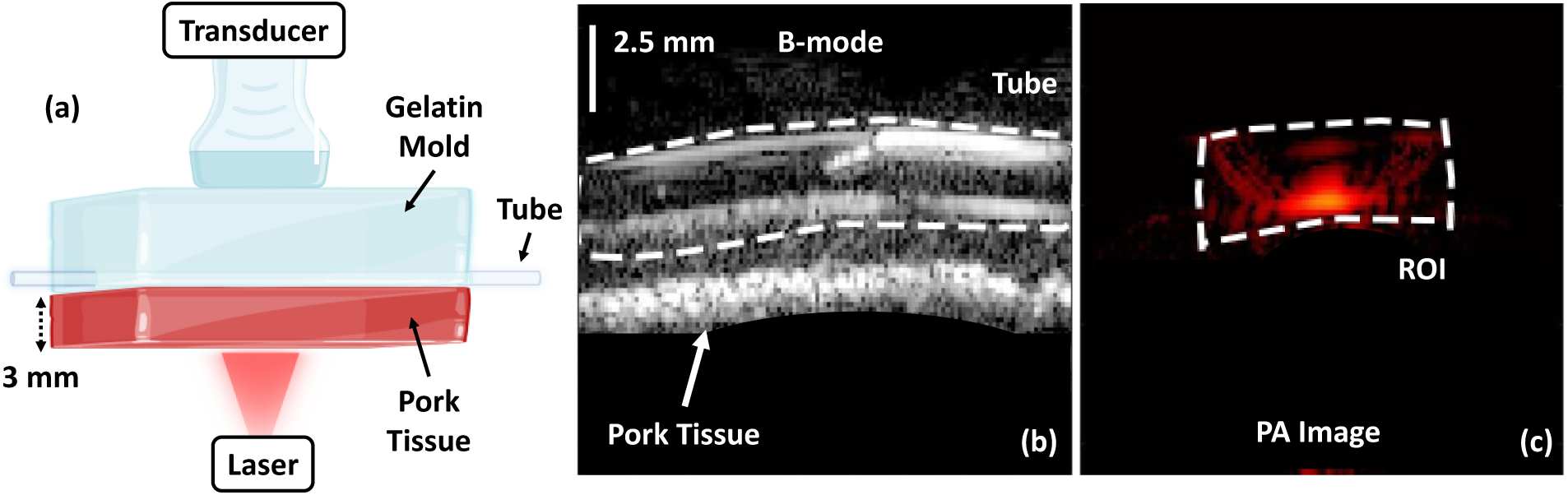
Porcine tissue phantom for NIR-II characterization of oxy-Hb and deoxy-Hb. (a) A schematic representation of the porcine tissue phantom used to characterize the NIR-II PA spectra of oxy-Hb and deoxy-Hb. (b) B-mode image of 3 mm slice of pork with a tube carrying blood underneath embedded in gelatin (scale bar = 2.5 mm). (c) PA image of tube carrying oxy-Hb collected at 1064 nm.

**Fig. 2:**
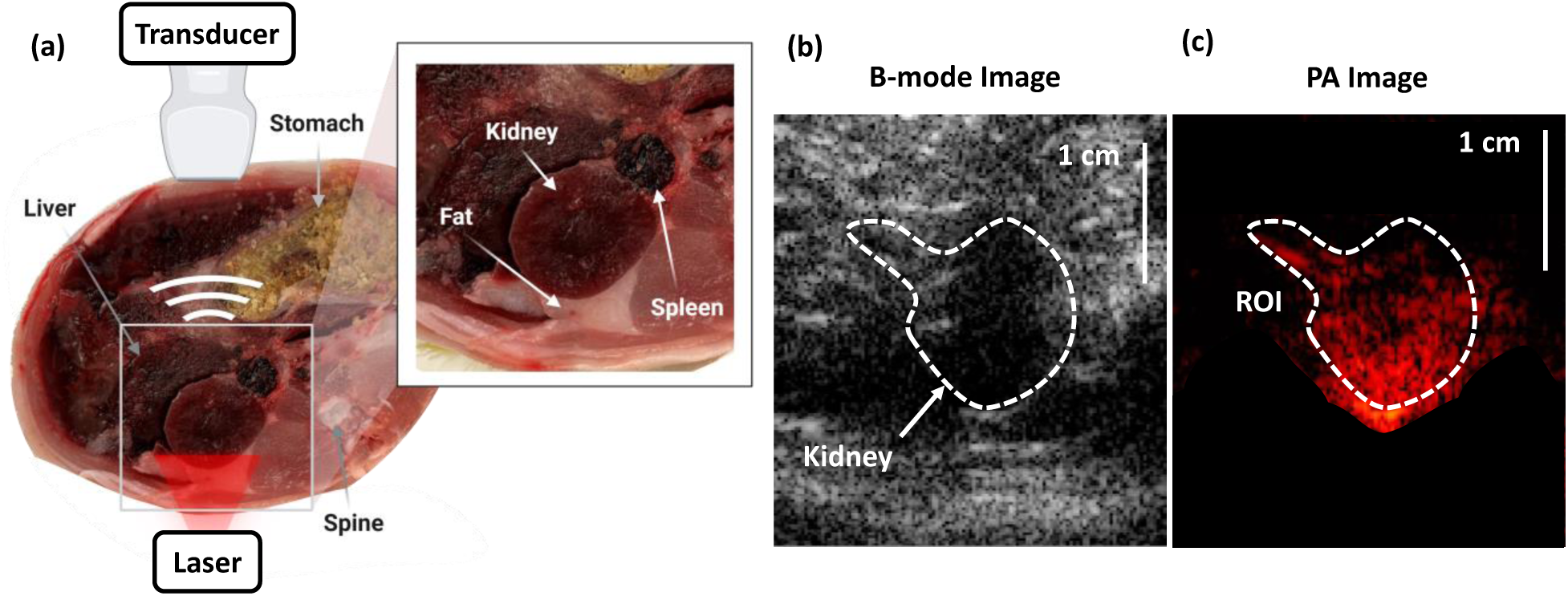
In vivo NIR-II PA imaging of rat kidney. (a) A photograph of a cross-section of the female rat abdomen with a description of the orientations of the light delivery and the acoustic detection for the renal imaging. The inset replicates the 2D slice of the rat abdomen acquired with the PA system (b) A B-mode image of the abdomen with a cross-section of the kidney (indicated with arrow). (c) PA image of the rat kidney collected at 1064 nm. The dashed white line represents the kidney region of interest.

### 2.2 Spectral Unmixing of Blood Oxygenation

The averaged PA signal (*p*_0_(λ)) were linearly unmixed to estimate the blood oxygenation (*sO*_2_) of imaged target using eq. 1.

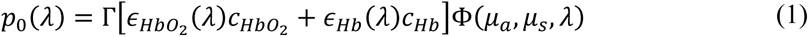

Here, ε_i_ is the molar optical extinction coefficient of the target chromophore – in this case, either oxygenated or deoxygenated hemoglobin, as a function of the illumination wavelength (λ), and ϕ represents the local optical fluence which is a function of both the wavelength and the optical properties of the ambient medium. *c_i_* is the distribution of the target chromophores (oxy-Hb & deoxy-Hb). Once the chromophore distributions are estimated, blood oxygenation can be calculated using eq. 2.

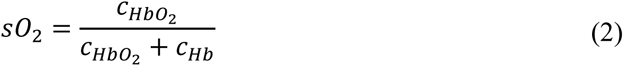

## 3. Results

### 3.1 NIR-II PA characterization of blood oxygenation

A porcine tissue phantom was used to characterize the PA spectra of whole oxy-Hb and deoxy-Hb blood over a wide spectral range. Figure 3a shows the PA spectra of three target solutions (oxy-Hb, deoxy-Hb and PBS) from 690 to 1400 nm (an extended spectral range up to 1800 nm is included in supplemental figure S3). In general, the fluence-normalized PA spectra of oxy-Hb and deoxy-Hb was consistent with the spectral shape of the optical absorption spectra (retrieved from literature [27]). Beyond 1300 nm, oxy-Hb and PBS are indistinguishable, likely due to the similar optical absorption of these species in this wavelength range. Beyond 1300 nm there is also an exponential rise in the optical absorption of water (which comprises 70% of all biological tissues), with diminishing differences in optical absorption between the three species (Fig. S3). However, we noticed a distinction between the three species at 1064 nm and a noticeable difference at 1200- 1300 nm, indicating that the use of wavelengths between 950-1300 nm may be sufficient for quantitative PA measurements.

**Fig. 3:**
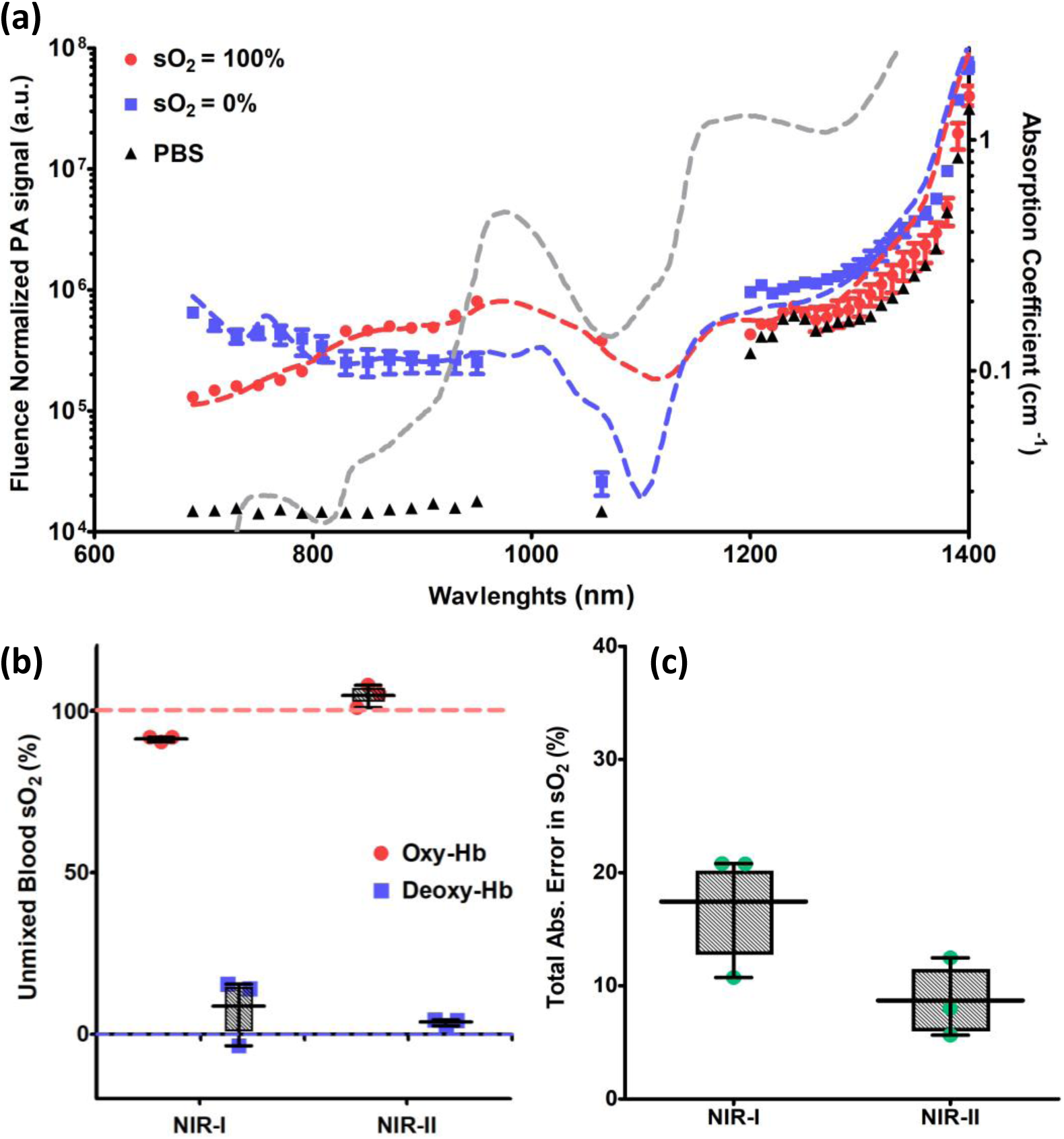
Porcine phantom imaging of whole blood. (a) A plot of local fluence normalized PA spectra from oxy-Hb (red circles), deoxy-Hb (blue squares) and PBS (black triangles) from NIR-I and NIR-II windows. Dashed lines represent the literature values (derived from [27]) of optical absorption of oxy-Hb (red), deoxy-Hb (blue) and water (gray) within this spectral window. (b) A box plot of spectrally unmixed blood sO_2_ using a combination of NIR-I (690, 808 & 950 nm) and NIR-II (1064, 1200 & 1230 nm) wavelengths. Colored dashed lines represent the true sO_2_ of the blood in the tube. (c) A boxplot of the total absolute error (linear sum of absolute errors in oxy-Hb and deoxy-Hb) in estimated sO_2_ using NIR-I and NIR-II wavelengths for all three phantom imaging runs. For all boxplots, the upper and lower limits of each box correspond to the 75th and 25th quartiles. The line through the center corresponds to the mean while the whiskers correspond to the range of the data.

Spectral unmixing to estimate blood oxygenation was performed using different combinations of wavelengths. Figure 3b shows a bar chart of the blood sO_2_, estimated using spectral unmixing, for two different combinations of laser wavelengths – NIR-I (690, 808 and 950 nm) and NIR-II (1064, 1200, 1230 nm). The total absolute error in the sO_2_ estimation (i.e. a sum of absolute error in estimated oxy-Hb & deoxy-Hb vs probe measurements) using these wavelength combinations is shown in figure 3c. The traditional NIR-I wavelengths used for spectral unmixing had the greatest error (∼15-20%), while NIR-II wavelengths had absolute errors under 10%.

### 3.2 In vivo validation of improved sPAI using NIR-II excitations

To validate the phantom experimental results, sPAI of in vivo renal sO_2_ was performed using wavelengths from the NIR-I and NIR-II windows. Figure 4a shows the absorption normalized PA spectra of rat kidneys under the three unique conditions (pre- & post-euthanasia, and bleached of hemoglobin) within the 690 to 1300 nm spectral range. We note that the normalized spectra of the PA signal and the optical absorption spectra show consistent trends between the different hemoglobin species. The PA spectra from oxy-Hb and deoxy-Hb kidneys were distinctly higher than kidney bleached of hemoglobin, confirming that the measured PA signals are generated by hemoglobin in the kidney. The highest contrast between oxy-Hb and deoxy-Hb (∼ 2× difference in CNR) is at 1064 nm. Furthermore, by imaging at the increased fluence available with NIR-II wavelengths, a 40% increase in the signal to noise ratio of deeply embedded vasculature was observed (Fig. S4).

**Fig. 4:**
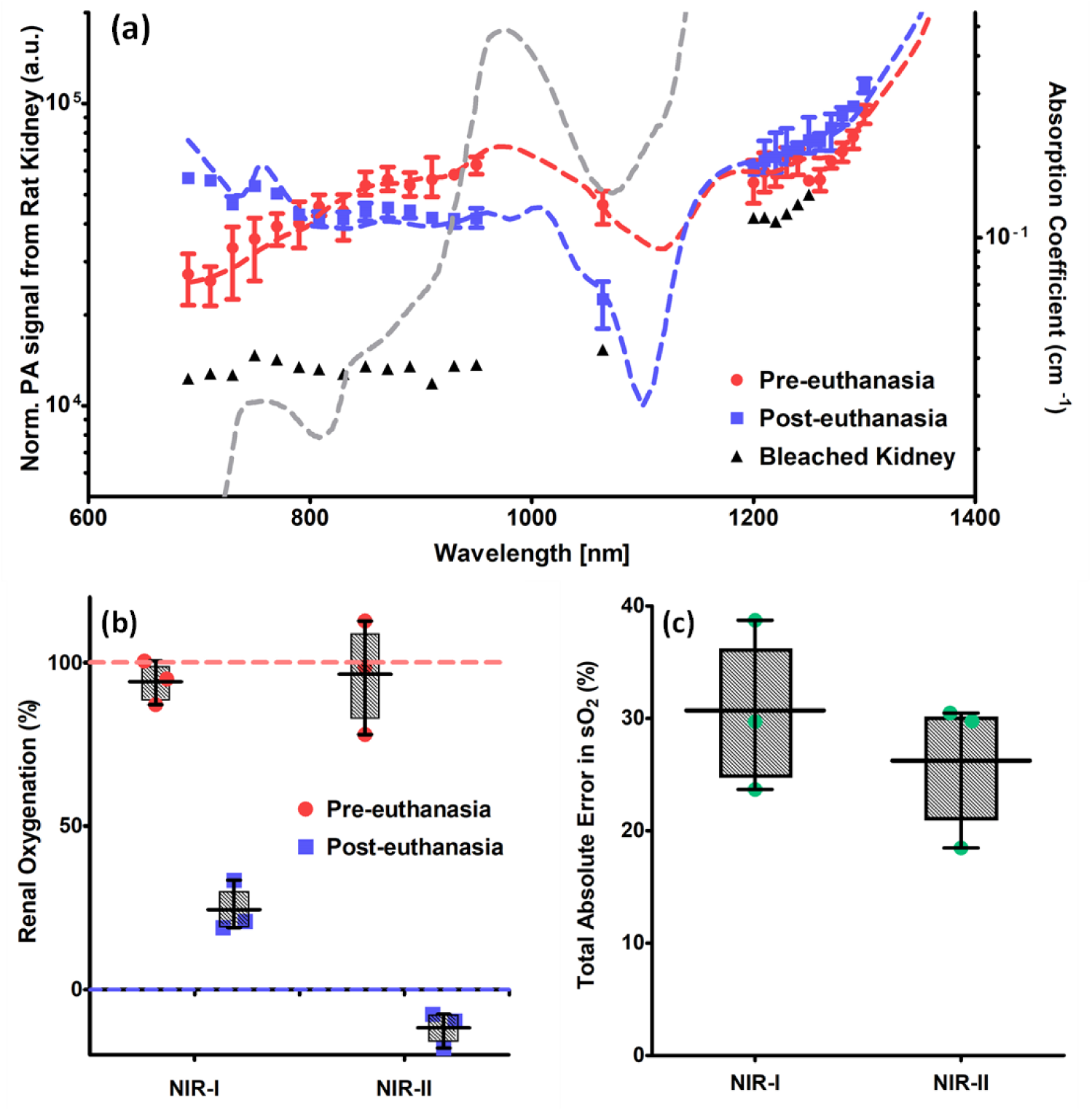
Spectral PA imaging of renal oxygenation using NIR-II wavelengths. (a) A plot of absorption normalized PA spectra of rat kidney imaged pre- (red) and post- (blue) euthanasia along with a rat kidney bleached of hemoglobin (black). The optical absorption spectra of oxy-Hb (red dashed lines), deoxy-Hb (blue dashed lines) and water (black dashed lines) is overlayed on the plot. (b) A plot of renal oxygenation estimated using spectral unmixing with combinations of wavelengths – NIR-I (690, 808 & 950 nm) and NIR-II (1064, 1200-1300 nm). Each box represents the unmixing results from segmented rat kidney (n = 3) before (red boxes) and after (blue boxes) euthanasia. The true renal oxygenation measured with an oxygen probe is indicated in dashed lines. (c) A plot of total absolute error in renal oxygenation (sum of absolute error in pre and post euthanized animal) for the two combinations of wavelengths used for spectral unmixing.

Renal sO_2_ was calculated using both the NIR-I and NIR-II wavelength ranges. Figure 4b shows a boxplot of the renal sO_2_ obtained using NIR-I (690, 808 and 950 nm) and NIR-II (1064, 1200- 1300 nm). Consistent with the phantom experiments, the PA spectral features follow the optical absorption spectra of each hemoglobin species. Oxygenation of the oxygenated and deoxygenated kidneys were measured with the oxygen probe to be ∼90% and 0%, respectively. The total absolute error in renal sO_2_ estimation (i.e. a sum of absolute error in sPA-derived renal sO_2_ compared to the oxygen probe measurements) is shown in figure 4c. There was an average spectral unmixing error of ∼30% when using traditional NIR-I wavelengths compared to ∼20% when imaging with NIR-II wavelengths.

## 4. Discussions

The primary objective of this work was to demonstrate spectral PA imaging of blood sO_2_ using NIR-II wavelengths. To achieve this, characterization of the PA spectra of oxy-Hb and deoxy-Hb blood was performed using a phantom (at a wide spectral range 690 – 1800 nm). The accuracy of spectral unmixing to estimate blood sO_2_ was assessed, and performance was validated *in vivo* in the kidneys of adult female rats. Two unique advantages facilitate deep tissue sPAI with the use of NIR-II wavelengths – the increased maximum permissible exposure limit (MPE) for skin, and reduced tissue scattering in this wavelength range.

Increased MPE limits in the NIR-II allow higher fluences to be delivered to the imaged target, thereby increasing the imaging depth and image contrast. Recently, Srishti et. al. demonstrated a 100% increase in imaging depth and a 30% increase in signal-to-noise ratio when using 1064 nm over NIR-I laser wavelengths using a simple ink-phantom [29]. They attribute these observations to the 5× increase in MPE (100 mJ/cm^2^) for NIR-II wavelengths versus NIR-I wavelengths (20 mJ/cm^2^ at 500 nm). Our experiments support these conclusions by generating visible PA contrast from the entire kidney and demonstrating a ∼40% increase in the signal to noise ratio of deeply embedded vasculature (such as the renal artery) when imaging using 1064 nm at ∼36% MPE (36 mJ/cm^2^) (Supplemental Fig. S4).

Reduced tissue scattering in the NIR-II also leads to improved light penetration and visualization of deeper structures. Tissue scattering remains a key limitation in light penetration depth. Various studies have demonstrated the improvement in photon penetration with the use of optical wavelengths that minimize tissue scattering for different imaging modalities [30–33]. Zhang et. al. demonstrated high tissue transparency within NIR-II spectral windows using hyperspectral optical imaging allowing for improved photon penetration and spatial contrast [30]. The use of NIR-II wavelengths for fluorescence imaging is a research area currently undergoing rapid expansion, with reduced tissue scattering facilitating whole body imaging in mice [32]. Using fluorescence imaging, Byrd et. al. demonstrated a 50% increase in the ratio of vessel-to-tissue fluorescence signal collected at NIR-II over NIR-I wavelengths, from an adult pig brain [31]. Similarly, deep tissue anatomical PA imaging using NIR-II wavelengths has been demonstrated [4, 29, 34, 35]. Recently, Na et. al. demonstrated high light penetration (up to 4 cm) using 1064 nm illumination allowing PA imaging of brain vasculature in humans [4]. Similarly, Schoustra et. al. demonstrated breast imaging using 1064 nm up to depths of 2.2 cm from the surface of the skin, using NIR-II wavelengths [36]. However, these studies were limited to single-wavelength anatomical imaging with no quantitative estimations of blood oxygenation through sPAI. Hence, to date, a systematic comparison of NIR-I against NIR-II excitations for sPAI of blood oxygenation has yet to be performed. In our studies, a ∼2× increase in light penetration both in phantoms and in vivo when using NIR-II wavelengths was observed (Fig. 3a & 4a).

Our results show improved accuracy in the measurement of blood and tissue oxygenation with the use of NIR-II compared to traditional NIR-I wavelengths. This improved accuracy of NIR-II may be attributed to reduced spectral coloring because of minimized tissue scattering (particularly between 1200-1300 nm). Recently, Hochuli et. al. showed that simple spectral unmixing using wavelengths in the NIR-I window yields highly inaccurate estimates of tissue oxygenation due to spectral coloring. They suggest the use of wavelengths where spectral coloring can be minimized while maintaining a sufficiently high signal-to-noise ratio [37]. In our phantom experiments, pork tissue has a total optical attenuation (absorption and scattering of water and lipids) of ∼50 cm^-1^ in the NIR-I window, dominated by diffused photon scattering. In contrast, tissue attenuation in the NIR-II window remains under ∼10 cm^-1^ (a 5× decrease), which is absorption-dominated. The Beer’s law based fluence compensation accurately calculates local fluence for deep targets in the presence of absorption dominated photon propagation, i.e. in the NIR-II window (as seen by a < 10% deviation between MC simulated and Beer’s Law calculated local fluences between 1200- 1300 nm is demonstrated in Fig. S5). Hence, NIR-II wavelengths offer improved sPAI of blood oxygenation from deeply embedded targets.

## 5. Conclusions

Our experimental results demonstrate sPAI imaging of tissue oxygenation using NIR-II wavelengths. We demonstrated improvement in the accuracy of spectral unmixing using NIR-II wavelengths in ex vivo porcine tissue phantoms and in vivo renal imaging. We attribute this improvement to the reduced spectral coloring due to reduced multiple tissue scattering and preferentially increased absorption of the imaged target in the NIR-II. We demonstrated these results up to a depth of ∼ 1.5 cm in vivo at a low fluence of 5-6 mJ/cm^2^ (5% of MPE). Future studies will explore the maximum possible imaging depth in larger animal models and at higher fluences.

## Acknowledgements

We thank Li Guang and Dr. Jonathon Quincy Brown for their recommendations on the tissue clearing protocol incorporated in this study. We thank Dr. Mistina Mano Manoharan for her editorial support.

## Supplemental Figures

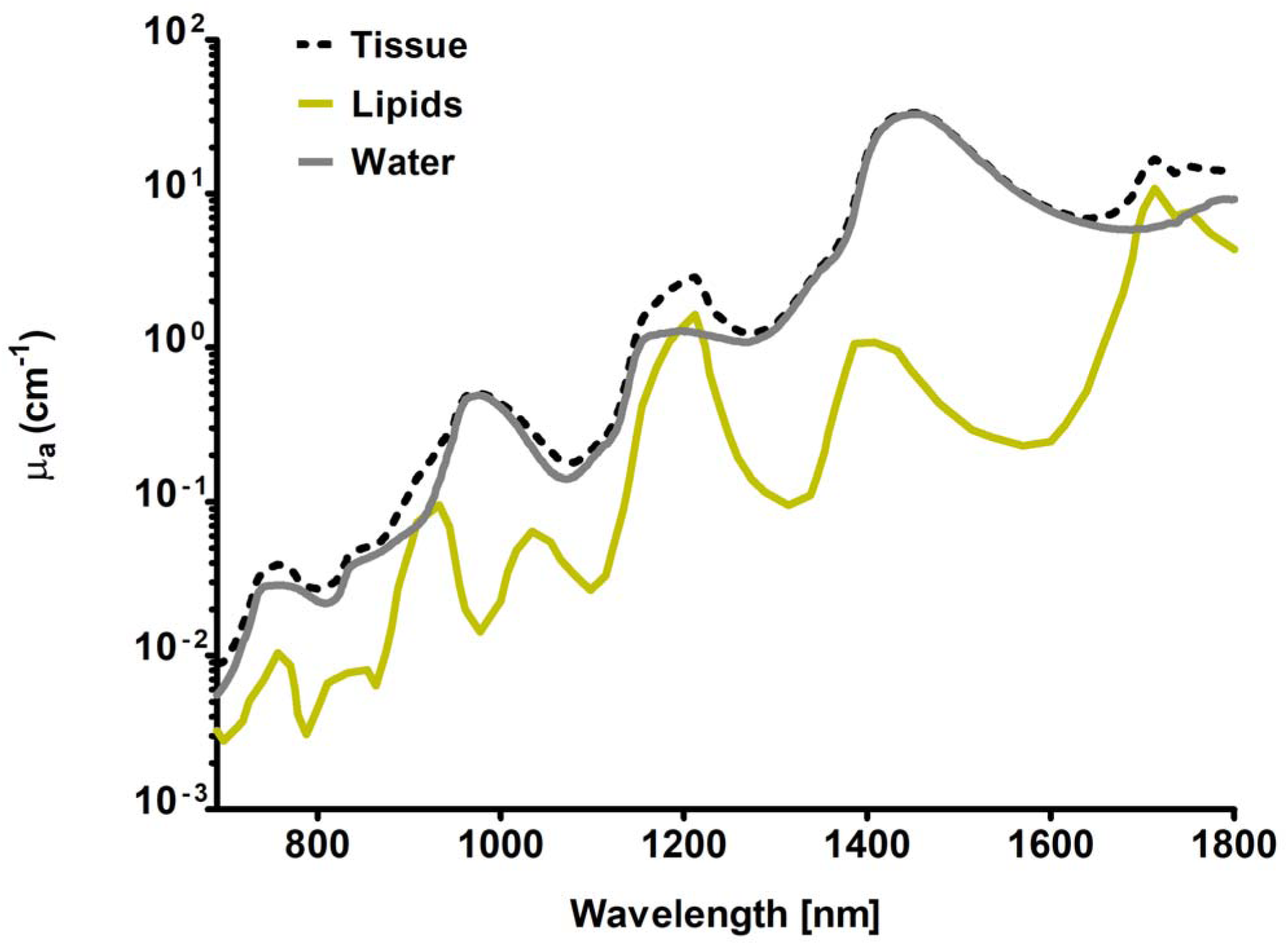

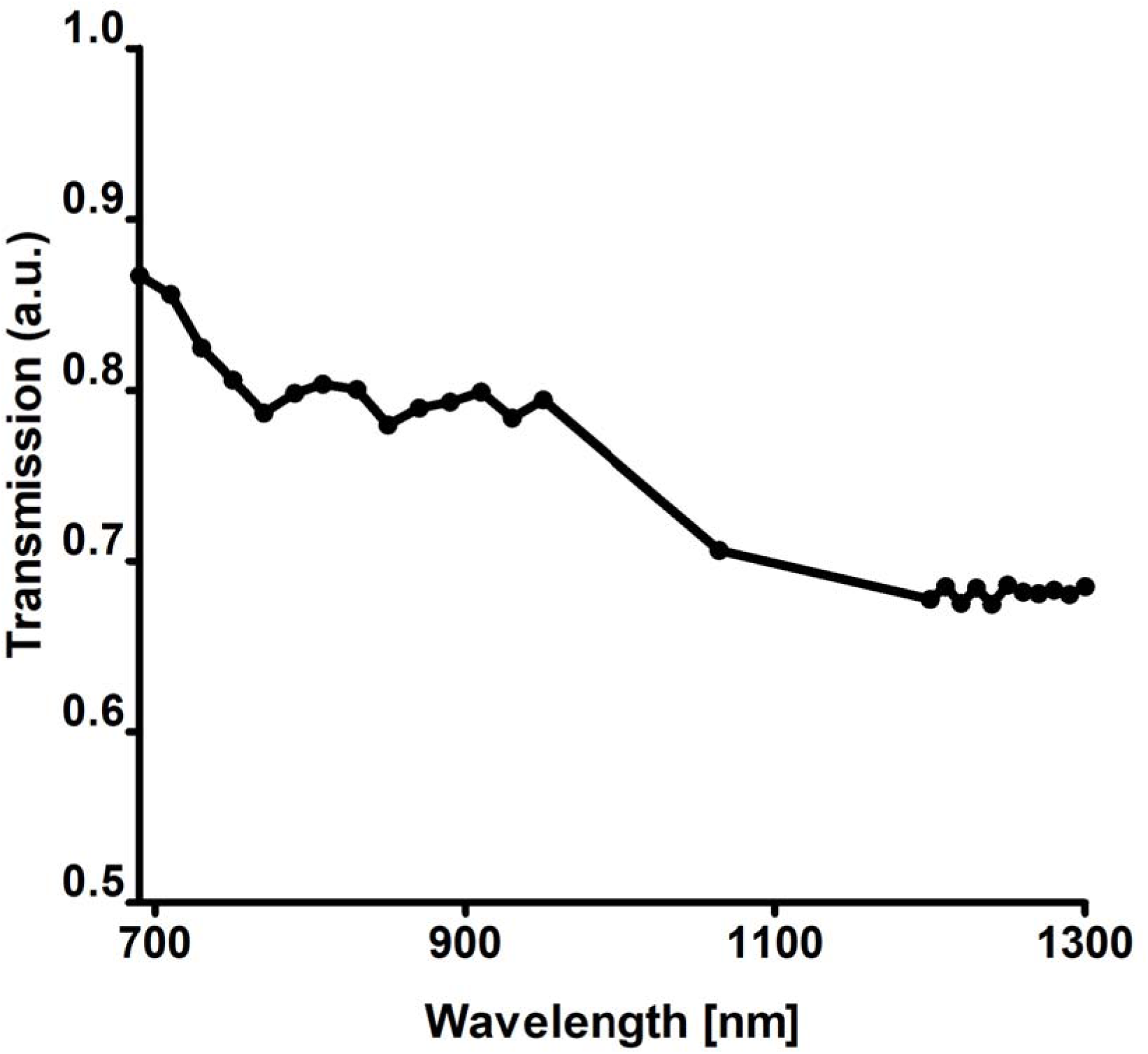

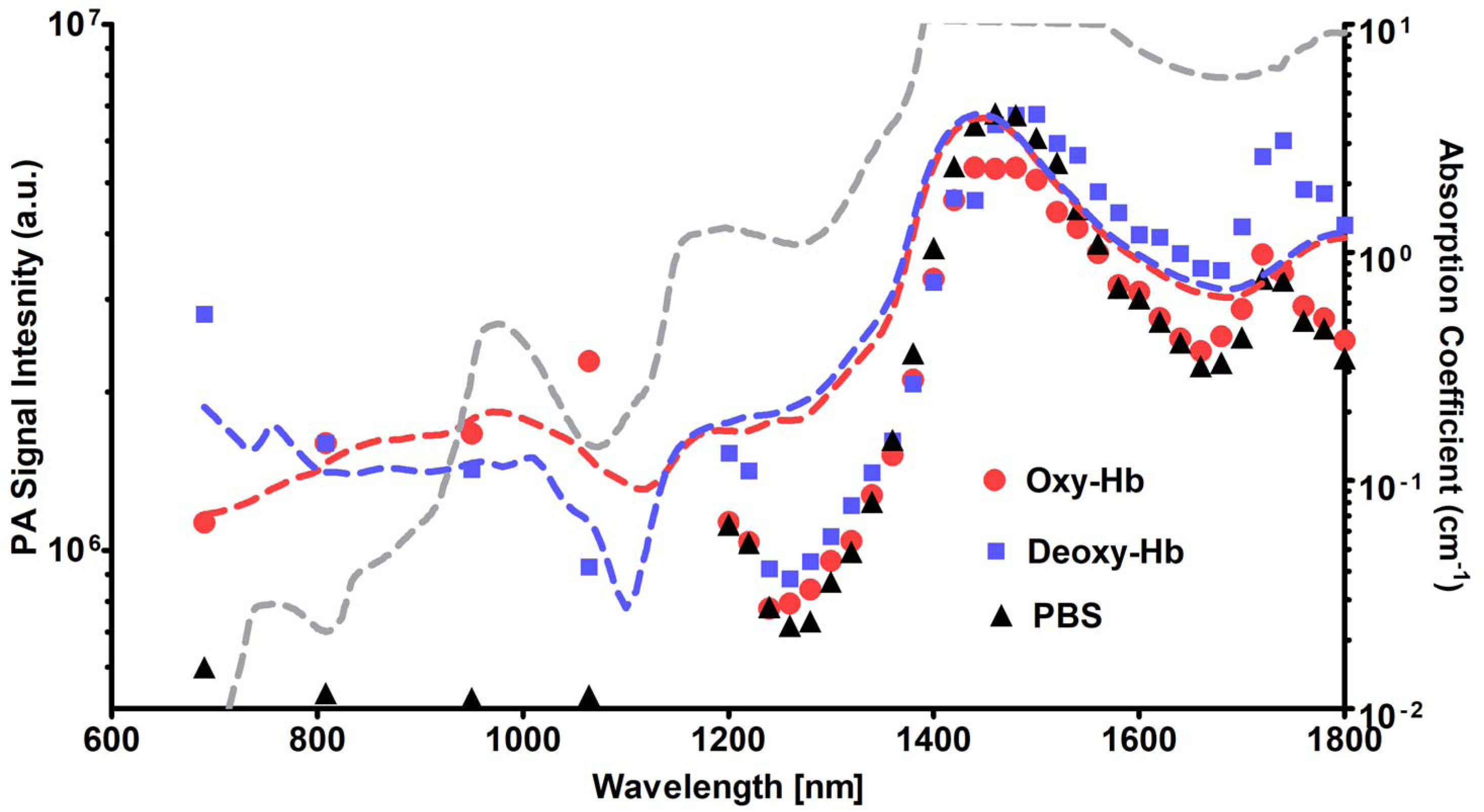

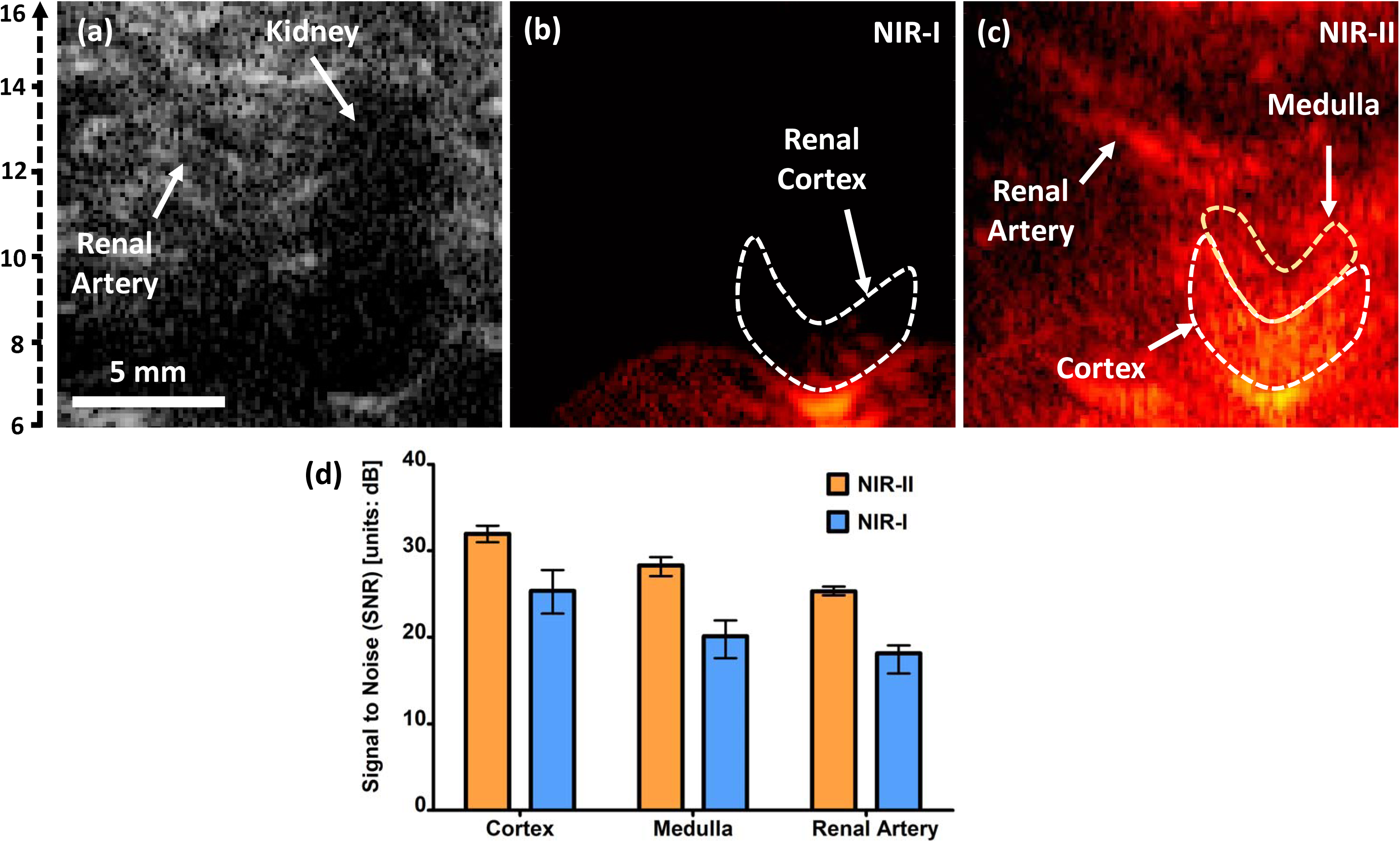

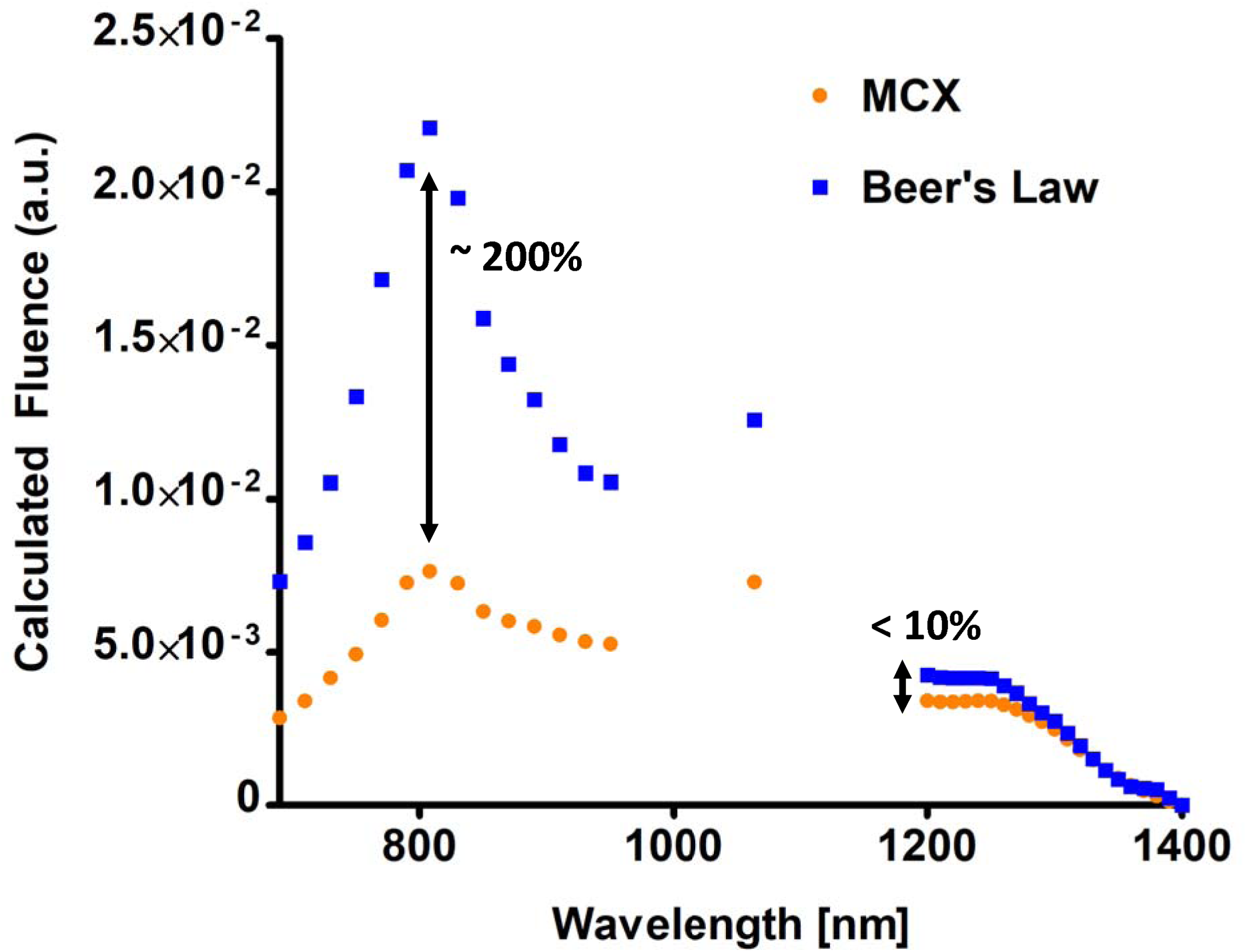

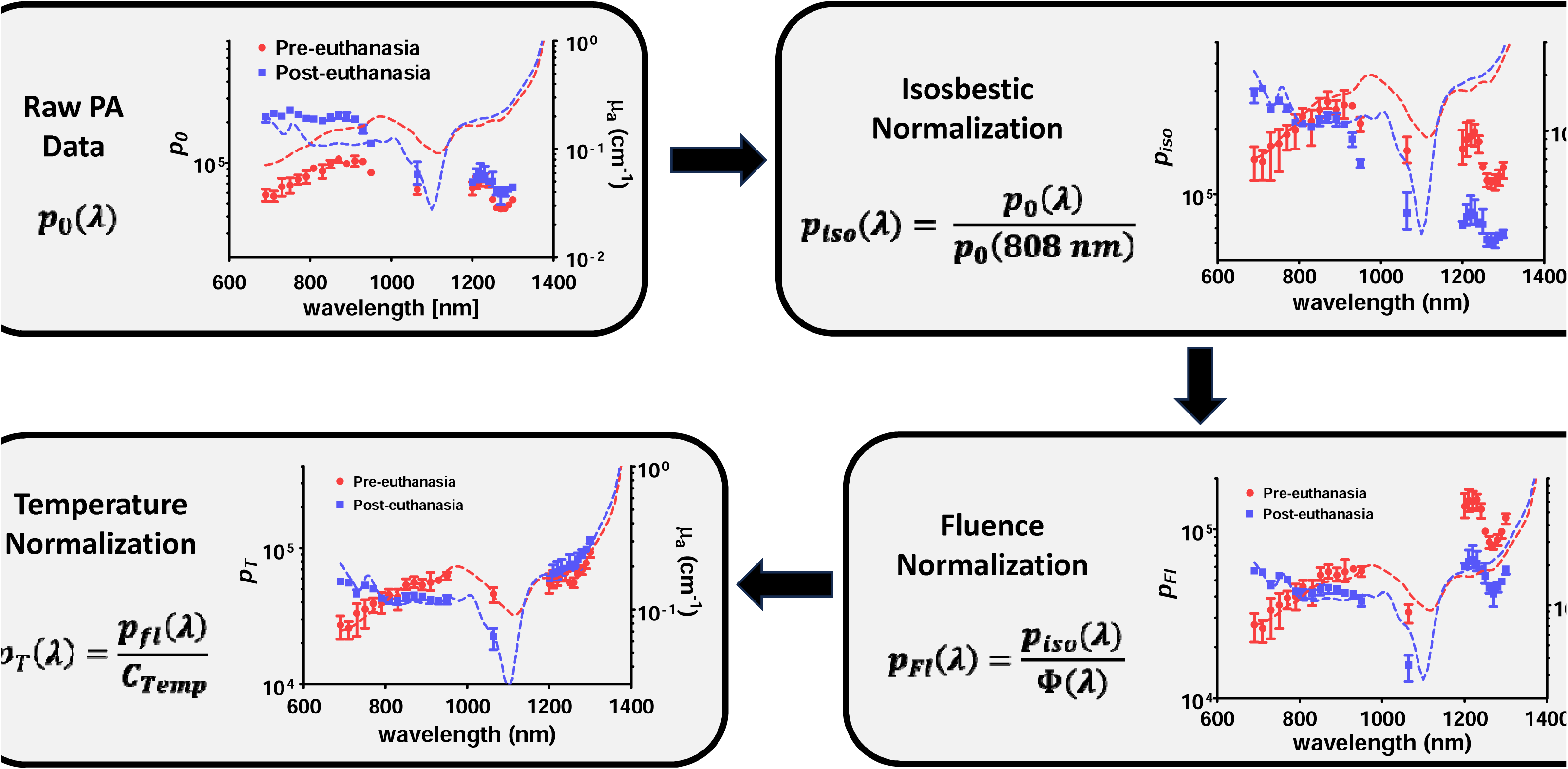

