## supplemental document for "Improved Spectral Inversion of Blood Oxygenation due to Reduced Tissue Scattering: Towards NIR-II Photoacoustic Imaging"

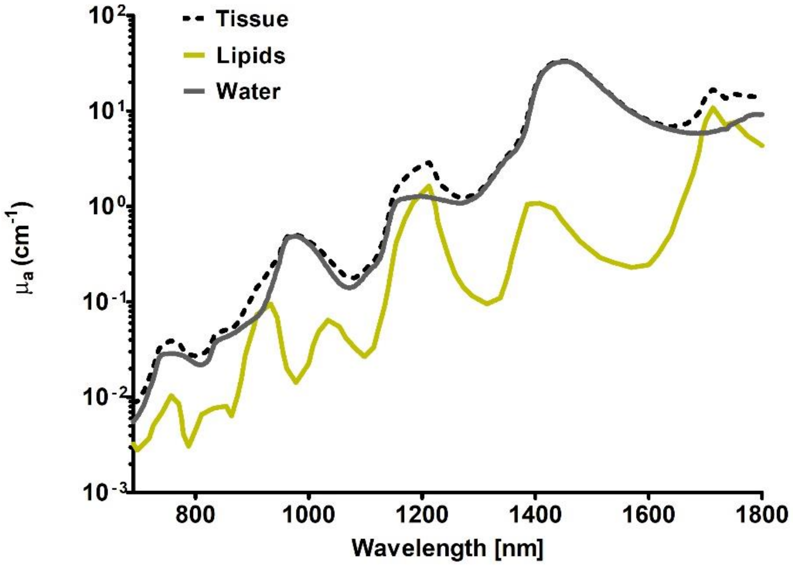


**Figure S1: Tissue attenuation used for fluence compensation**. A plot of the tissue absorption coefficients (adapted from [1]) used to compensate local fluence of the photoacoustic images.

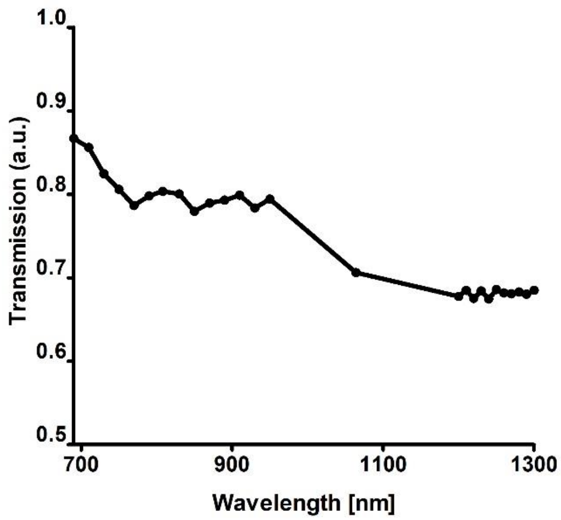


**Figure S2: Light attenuation by the transparent (polyethylene terephthalate, PET) sheet**. A plot of the transmission of light through the PET sheet used for in vivo rat imaging.

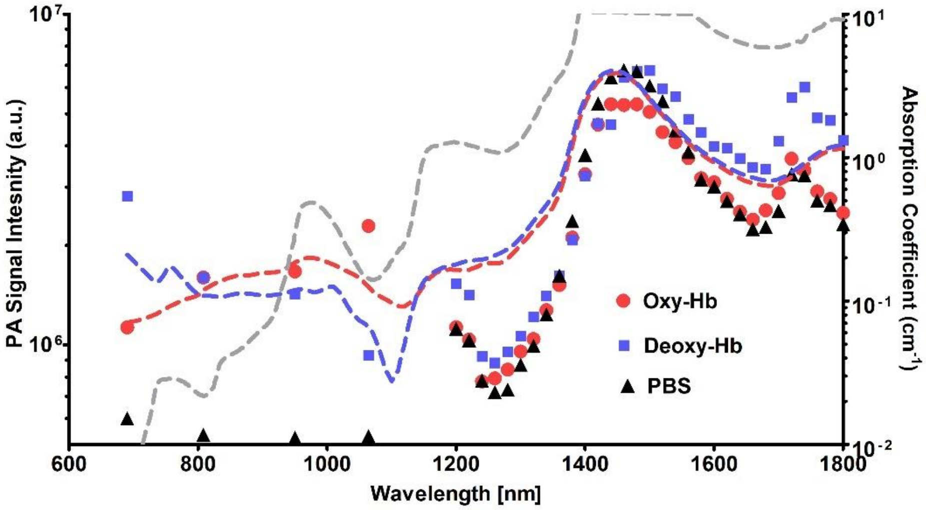


**Figure S3: Extended PA spectra of whole blood**. PA spectra of whole blood up to 1800 nm – oxygenated (red circles) and deoxygenated (blue squares) and PBS (black triangles) measured in a porcine phantom.

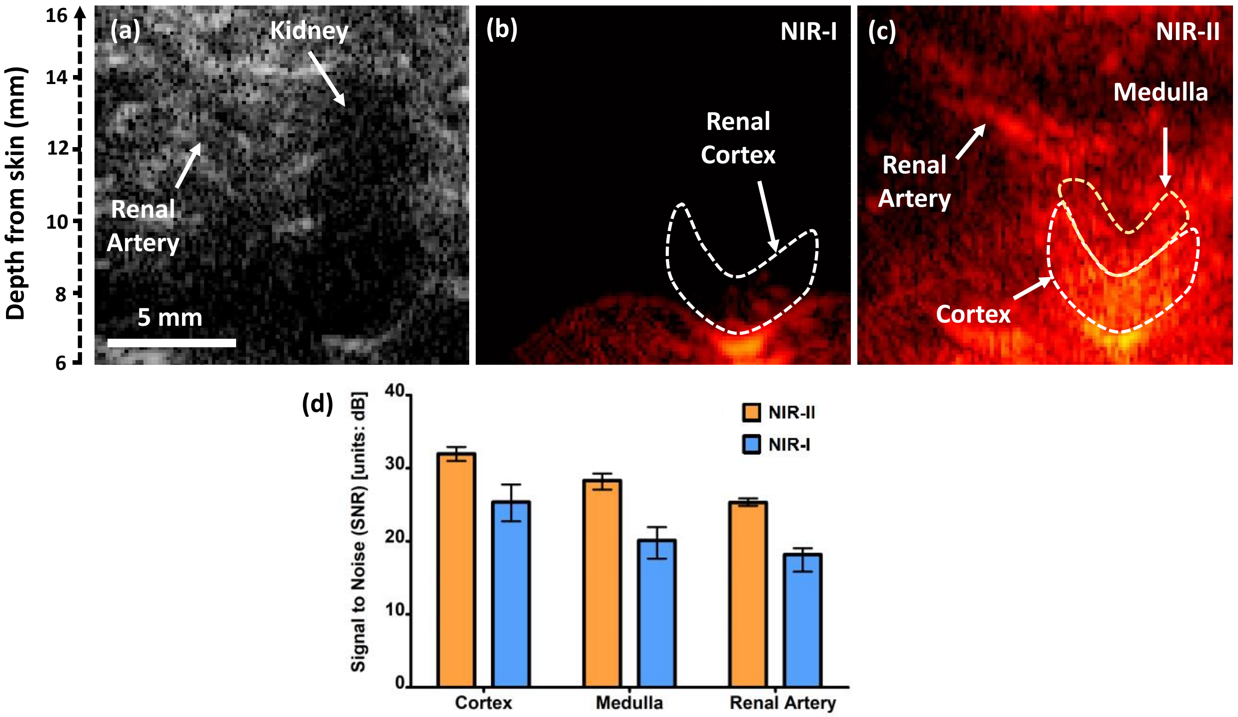


**Figure S4: Improvement in signal to noise ratio (SNR) with NIR-II at higher fluences.** B-mode (a) and PA images (b-c) of oxygenated rat kidney acquired at 690 nm (NIR-I, Fig. S4b) and 1064 nm (NIR-II, Fig. S4c) with optical fluence at the surface of skin of 6 mJ/cm$^2$ (b) and 36 mJ/cm$^2$ (c), respectively. The imaging depth, relative to the skin surface, is shown as the y-axis in (a-c). (d) A plot of the signal to noise (SNR) ratio (units of dB) of the different anatomical regions. Orange and blue bars correspond to mean SNR at NIR-II and NIR-I, respectively, with error bars representing the standard deviation of five acquisitions.

Monte Carlo simulations were performed using the MCX toolbox [2]. Here, a simple homogenous slab of media with dimensions of 1 x 1 x 1.2 cm and properties set porcine tissue, was defined (retrieved from [3]). A cylindrical inclusion (diameter of 1.2 mm) with properties of whole oxygenated blood was defined at a depth of 3 mm from the surface of illumination. A gaussian beam of 0.5 cm diameter was centered on the sample at the illumination surface. Simulations were performed in the NIR-I (690 – 950 nm in 20 nm increments) and NIR-II (1200 – 1400 nm in 10 nm increments). The fluence deposited at the tube was averaged and used for further analyses. The local fluence at the cylindrical inclusion was also calculated using Beer's law which can be represented using the following equation,

$$\Phi(z,\lambda) = \Phi_0 e^{-\mu_a(\lambda)\, z} \tag{1}$$

where $\Phi(z,\lambda)$ is the fluence at a depth of $z$ within the simulated volume and at wavelength $\lambda$. $\Phi_0$ is the fluence at the illumination surface while $\mu_a(\lambda)$ is the absorption coefficient of pork muscle at the respective wavelength.

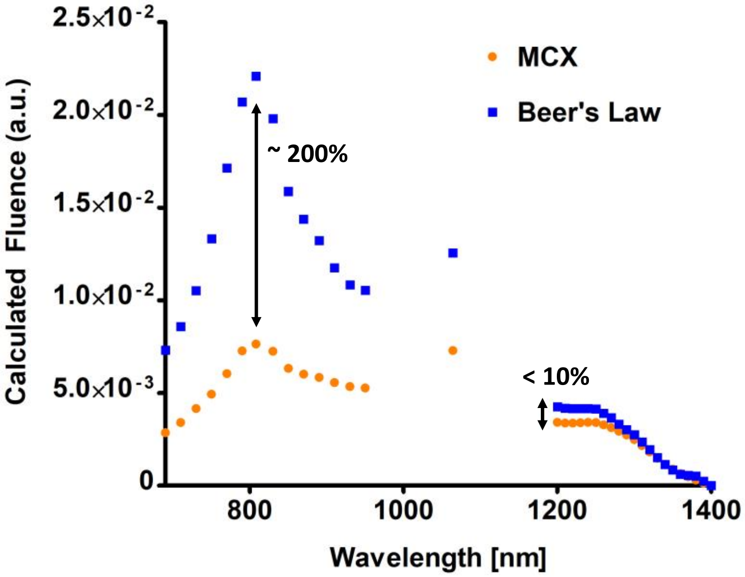


**Figure S5: The effects of scattering on local fluence in NIR-I vs NIR-II.** A plot showing the local fluence simulated using MC modelling and using Beer's law. The average percent difference between the two spectra in the NIR-I and NIR-II regions are considerably different, and this may contribute to more accurate quantification of oxygen saturation in the NIR-II.

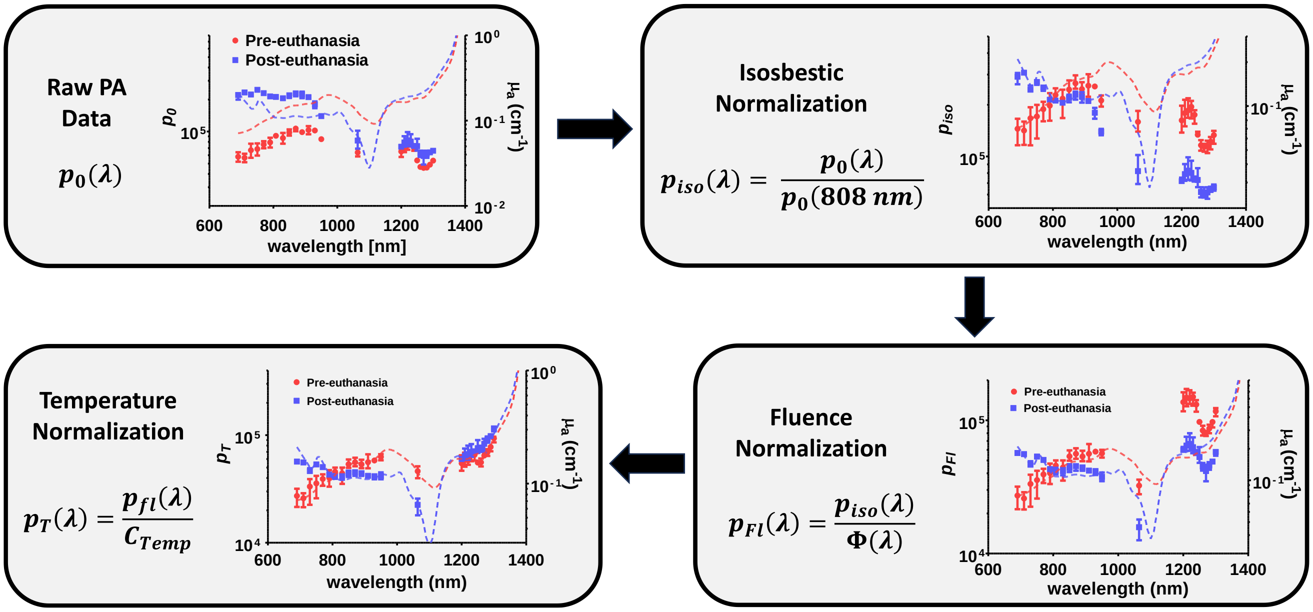


**Figure S6: Data correction scheme of in vivo data.** A flowchart describing the correction of the raw PA data collected from in vivo rat kidney. The correction includes a division of the pre- and post-euthanized PA spectra at to match at the isobestic wavelength (808 nm). This is followed by a division of both spectra by the wavelength dependent local fluence. Finally, the PA spectra beyond 1200 nm of pre-euthanized rat kidney was divided by a factor ($C_{Temp}$) that is ratio of pre- to post- animal temperature
